# *Theileria* highjacks JNK2 into a complex with the macroschizont GPI-anchored surface protein p104

**DOI:** 10.1101/383646

**Authors:** Perle Latré De Laté, Malak Haidar, Hifzur Ansari, Shahin Tajeri, Eszter Szarka, Anita Alexa, Kerry Woods, Attila Reményi, Arnab Pain, Gordon Langsley

## Abstract

*Theileria* is a unique apicomplexan parasite capable of transforming its host cell into a disseminating tumour. Constitutive JNK activity characterizes bovine T and B cells infected with *T. parva*, and B cells and macrophages infected with *T. annulata.* Here, we show that *T. annulata* manipulates JNK activation by recruiting JNK2, and not JNK1, to the parasite surface, whereas JNK1 is found predominantly in the host cell nucleus. *In silico* analysis identified 3 potential JNK-binding motifs in the previously characterized GPI-anchored macroschizont surface protein (p104), and we demonstrate here that JNK2 is recruited to the parasite via physical interaction with p104. A cell penetrating peptide harbouring a p104 JNK-binding motif also conserved in *T. parva* p104 competitively ablated binding, whereupon liberated JNK2 became ubiquitinated and degraded. Sequestration of JNK2 depended on PKA-mediated phosphorylation of the conserved JNK-binding motif and upon disruption of the p104/JNK2 complex loss of JNK2 resulted in diminished matrigel traversal of *T. annulata*-transformed macrophages. Loss of JNK2 also resulted in upregulation of small mitochondrial ARF that promoted autophagy consistent with cytosolic sequestration of JNK2 sustaininf not only JNK2, but also nuclear JNK1 levels that combined contribute to both survival and dissemination of *Theileria*-transformed macrophages.

**Author Summary:** *Theileria annulata* parasites infect and transform their host bovine leukocytes into tumour-like cells that disseminate throughout infected animals causing a widespread disease called tropical theileriosis. Virulence has been ascribed to the parasite’s ability to constitutively activate leukocyte c-Jun N-terminal Kinase (JNK) leading to permanent induction of Matrix Metallo-Proteinase 9 (MMP9) that promotes transformed macrophage dissemination. In attenuated live vaccines JNK-mediated AP-1-driven transcriptional activity is reduced so dampening dissemination. However, in leukocytes JNK exists as two isoforms JNK1 and JNK2 and here, we show for the first time that in *T. annulata*-transformed macrophages they have different subcellular localisations and perform separate functions. Surprisingly, JNK2 associates with the parasite and is not in the nucleus like JNK1. JNK2 is hijacked by the parasite and sequestered in a complex with a macroschizont surface protein called p104. Upon forced complex dissociation JNK2 gets degraded and its loss negatively affects infected macrophage survival and ability to disseminate.

## Introduction

In mammals, c-Jun-N-terminal kinase (JNK) is encoded by three genes, *mapk8, mapk9* and *mapk10: mapk8* and *mapk9* code, respectively, for the ubiquitously expressed JNK1 and JNK2 proteins, and *mapk 10* codes for JNK3, whose expression is restricted to cardiac, nervous and testicular tissues [1]. JNKs are activated by environmental stress including extracellular insults such as radiation, redox, osmotic and temperature shocks, and intracellular stress such as miss-folded proteins [1]. Biological responses transduced through JNK-dependent pathways encompass proliferation, migration, survival, differentiation, inflammation [1] and some of the JNK substrates participating in these processes have been identified [2]. Changes in gene expression resulting from JNK activation may be accounted for by the phosphorylation of several transcription factors and the ensuing alteration in their transcriptional activity [2]. A well-characterized substrate of JNK is c-Jun, a component of the AP-1 transcription factor that is essential for proliferation and cell survival. JNK can affect c-Jun both positively and negatively: N-terminal phosphorylated c-Jun displays increased trans-activating activity [3], whereas in neurons the E3 ubiquitin ligase SCF^Fbw7^ specifically targets phosphorylated c-Jun for proteasome degradation, thereby controlling the JNK/c-Jun apoptotic pathway [4]. Additionally, in T lymphocytes c-Jun turnover is regulated by the E3 ubiquitin ligase Itch, whose activity increases upon JNK-dependent phosphorylation [5]. JNK can promote cell motility via alteration of focal adhesion dynamics following JNK-mediated phosphorylation of the focal adhesion adaptor paxillin [6-8].

Importantly, loss of *jnk2* in mouse embryonic fibroblasts (MEFs) increases cell proliferation, whereas a loss of *jnk1* leads to decreased proliferation and these contrasting effects are attributed to differential regulation of the mitogenic transcription factor c-Jun. JNK1 increases c-Jun stability via phosphorylation of serine 73, whereas JNK2 decreases c-Jun stability by promoting its ubiquitination [9, 10]. JNK2 also promotes ubiquitination-dependent proteasomal degradation of small mitochondrial ARF (smARF), as in *jnk*2^-/-^ MEFs (Mouse Embryonic Fibroblasts) levels of smARF rise to induce autophagy [11, 12]. SmARF is a short isoform of the tumour suppressor p19^ARF^ and interestingly, suppression of smARF did not require the kinase activity of JNK2 consistent with JNK2 acting as a scaffold protein [12]. The above examples highlight the disparate functions of JNK isoforms and underscore the necessity to study them individually and together to properly grasp the cellular impact of JNK activation.

Parasites of the genus *Theileria* are intracellular protozoans belonging to the *Apicomplexa* phylum. *T. annulata* and *T. parva* are two particularly pathogenic species that cause bovine lymphoproliferative diseases, respectively named tropical theileriosis and the East Coast Fever (ECF). The target host cells of *T. parva* are T-and B-lymphocytes, whereas monocytes/macrophages and B cells are preferentially infected by *T. annulata*. ECF and tropical theileriosis display similarities to human lymphomas and myeloid leukemias. A live attenuated vaccine exists to tropical theileriosis [13] that is generated by multiple passages of infected monocytes/macrophages, which become attenuated having lost their hyper-disseminating virulence trait [14]. *Theileria-*infected leukocytes behave as transformed cell lines, as they no longer require exogenous growth or survival factors, can form colonies in soft agar and give rise to disseminated tumours in immuno-compromised mice [15, 16]. Known as the only eukaryote pathogen to transform a eukaryote host cell *Theileria* achieves this by manipulating host cell signalling pathways, reviewed in [17]. Several different signalling pathways have been implicated, including TGF-β [18-20] and JNK kinase leading to constitutive phosphorylation of c-Jun and activation of AP-1 [14, 21-23].

Remarkably, *Theileria*-induced transformation is strictly dependent on the presence of live parasites, as the transformed host cell phenotype is fully reversible upon drug-induced parasite death; drug-treated transformed leukocytes return to a quiescent, non-activated state,and eventually die [24]. *Theileria*-dependent JNK1 activity is required for survival of *T. parva* transformed B lymphocytes, as demonstrated by over expression of a dominant negative kinase-dead mutant of JNK1 and/or via the use of pan JNK inhibitor [16, 25]. While the parasite-derived molecular mechanism(s) underlying JNK activation is/are unknown, JNK1-mediated survival of *Theileria*-transformed leukocytes has been attributed to AP-1-driven expression of the anti-apoptotic genes *Mcl-1* and *c-IAP* [16], and uncontrolled proliferation linked to AP-1-driven expression of transferrin receptor and cyclin D1 [23].

One of the characteristics of *Theileria*-transformed leukocytes is they display heightened oxidative stress due in part to uncontrolled proliferation-related production of H_2_O_2_ [26, 27]. This raises the possibility that exposure to H2O2 contributes to induction of JNK activity, as JNK activation occurs in response to stress. Taken together, JNK activation seems a key event in *Theileria*-induced leukocyte transformation and the aim of this study was to examine whether *Theileria* infection affects differentially JNK1 versus JNK2 and do the two isoforms play similar or different roles in parasite-induced leukocyte transformation.

## Results

### Cytosolic localization of JNK2 versus nuclear localization of JNK1 in *Theileria*-infected macrophages

To understand how the two major JNK isoforms participate in *Theileria*-induced leukocyte transformation we ascertained the sub-cellular distribution of JNK1 versus JNK2 in *Theileria-* infected macrophages. JNK1 partitions into the cytosolic and nuclear fractions and expression levels are parasite-dependent, decreasing upon Bw720c-induced parasite death, and this is particularly obvious for nuclear JNK1 (Figure 1A, left). In contrast to JNK1, JNK2 partitions principally in the cytosolic fraction and again levels decrease upon Bw720c (Fig 1B). Immunofluorescence analysis revealed JNK2 in the cytosol, decorating the intracellular macroschizont highlighted with a monoclonal antibody (1C12) to *T. annulata* p104 [28].

**Fig. 1.**
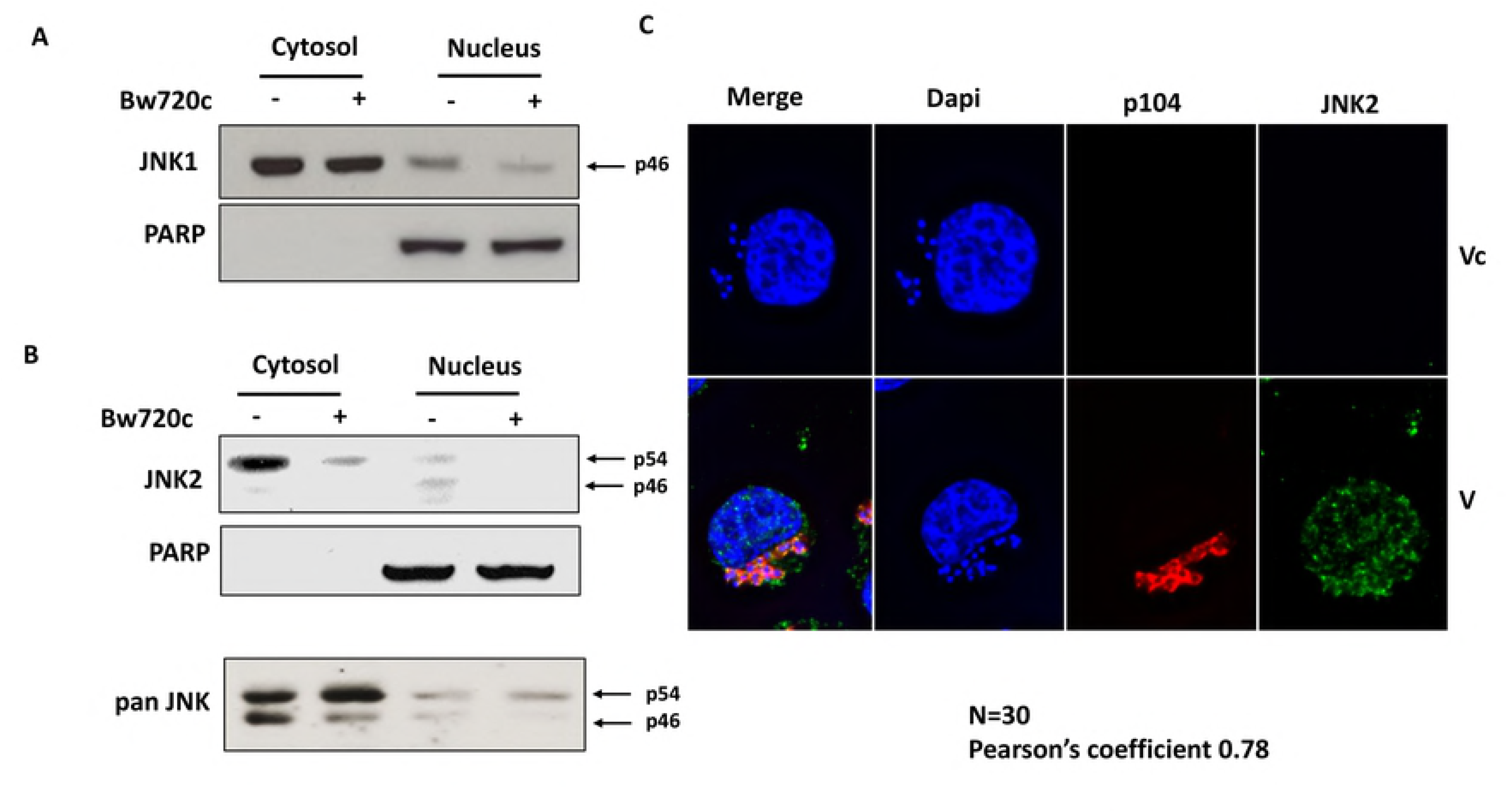
JNK2 is predominantly in the infected macrophage cytosol associated with the parasite. Localization of JNK1 (A) and JNK2 (B) in Theileria-infected macrophages treated or not with parasiticide drug BW720c. (A) Western blot analysis of nuclear and cytosolic extracts probed with a specific anti-JNK1 antibody and (B) Western blot of nuclear and cytosolic extracts probe with a specific anti-JNK2 antibody. (C) Immunofluorescence image showing association of JNK2 with the parasite decorated by a parasite monoclonal antibody (1C12) to P104. JNK2/P104 co-localization was analysed by the Manders method of pixel intensity correlation measurements using ImageJ/Fiji-Coloc2 plugin, and an average for 30 independent cells is given. DNA was stained with DAPI (blue).

### *T. annulata* p104 is a putative macroschizont surface JNK-binding protein

As JNK2 appears associated with the macroschizont we searched for a parasite surface protein predicted to have a JNK-binding motif [2]. *In silico* analyses were performed on three different species of *Theileria (T. annulata, T. parva* and *T. orientalis).* A Dfinder scan of the whole predicted proteomes of three *Theileria* species in search of D-motifs [29] led to identification of 26, 24 and 25 proteins, respectively (S1 file). Next, we asked which of these 75 putative *Theileria* JNK-binding proteins also had a predicted signal peptide and this criterion identified only one protein in *T. annulata* (TA08425), two proteins in *T. parva* (TP04_0437 orthologue of TA08425, and a hypothetical protein TP02_0553) and no protein in *T. orientalis.* TA08425 codes for a GPI-anchored *T. annulata* macroschizont surface protein called p104 [28] that has 3 putative decapeptide D-motifs KNESMLRLDL, KSPKRPESLD and KRSKSFDDLT located between amino acids 331-341, 606-616 and 804-814, respectively. However, only the decapeptide motif between amino acids 804 and 814 is conserved in the *T. parva* p104 orthologue (TP04_0437). The amino acid sequence in this region of p104 is not conserved in the non-transforming *T. orientalis* (TOT_040000478).

We examined therefore, whether the conserved D-motif (KRSKSFDDLT) mediated JNK2 binding to the *T. annulata* macroschizont surface protein p104. First, a pan-JNK antibody precipitated endogenous p104 from extracts of *T. annulata*-infected TBL3 B cells, but not from uninfected BL3 B cells (Figure 2A). As expected (see Fig 1), p104 was preferentially found in pan JNK precipitates (Fig 2B, left), and specifically in JNK2 precipitates (Fig 2B, right and Fig 3B). Altogether these results suggest that JNK2 is associated with p104 at the surface of the *Theileria* macroschizont.

**Fig. 2.**
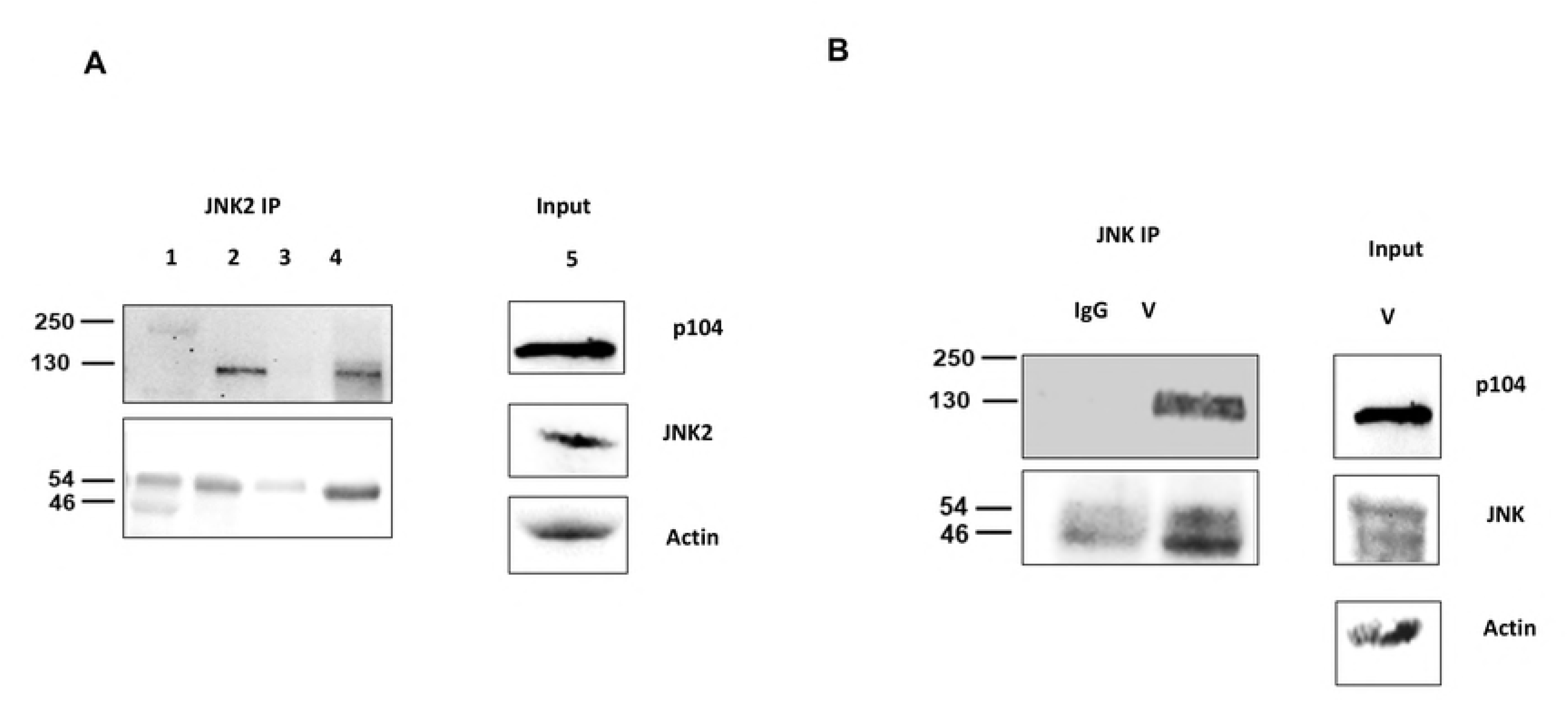
p104 interacts with JNK2 in *Theileria-infected* leukocytes. (A) Immunoprecipitation with a pan anti-JNK (JNK-IP) antibody using whole cell lysates derived from infected (TBL3) and non-infected B cells (BL3), with the precipitate probed with the anti-p104 monoclonal antibody 1C12. Input, shows JNK and P104 protein levels in BL3 and TBL3 cells revealed by respective antibodies. (1): BL3; (2): TBL3 1μM; (3): IgG; (4): V and (5): input form V. (B) Right, Immunoprecipitation with a pan-JNK, specific anti-JNK2, and irrelevant IgG control antibodies with the precipitate from infected macrophages (V) probed with 1C12. Input, shows (arrowed) the levels of the p46 JNK1 and p54 JNK2 isoforms revealed with the pan-JNK antibody. An anti-actin antibody was used as a loading control.

**Fig. 3.**
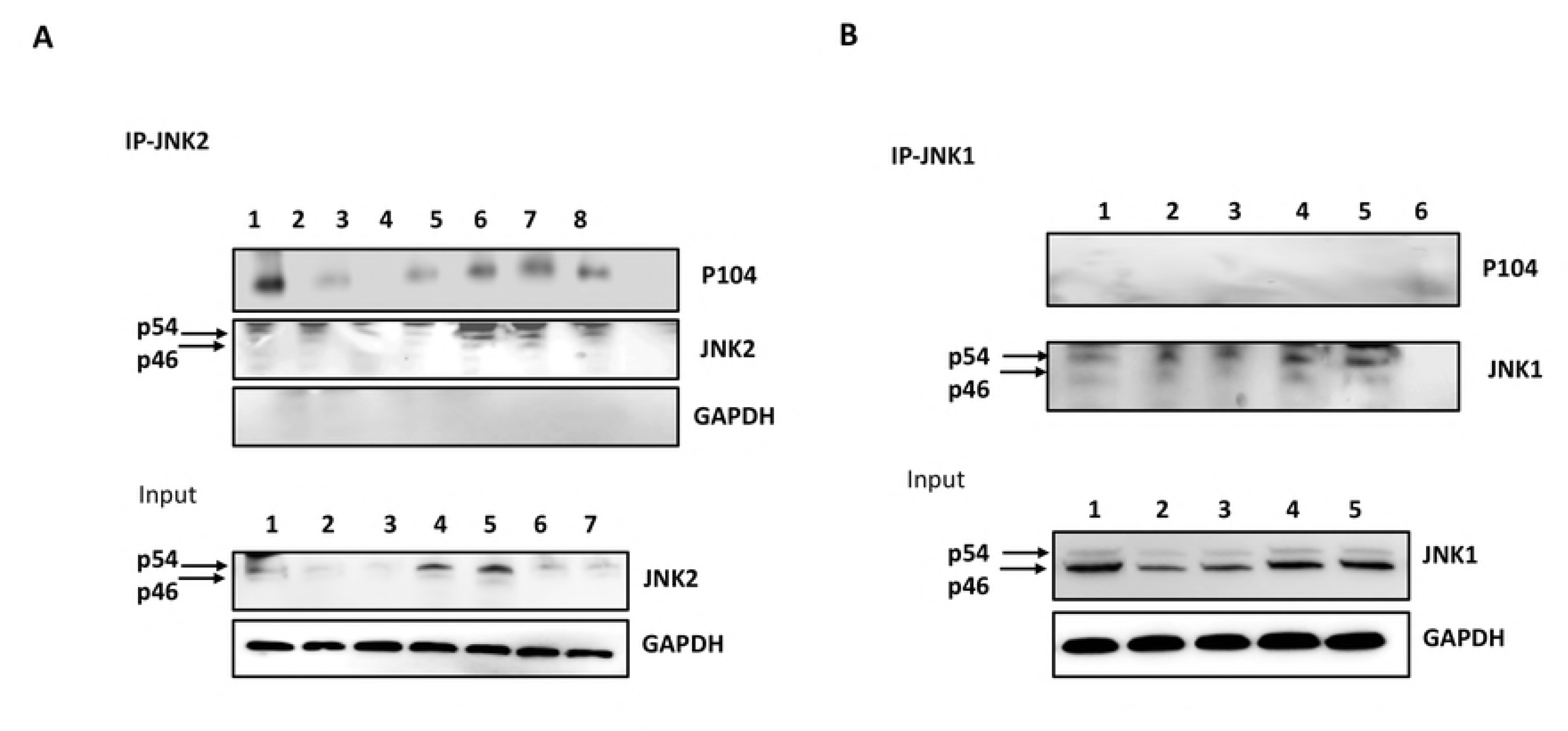
Abrogation of JNK2/p104 interaction leads to proteasome-mediated JNK2 degradation. Immunoprecipitation analyses with specific anti-JNK2 and anti-JNK1 antibodies using whole cell lysates derived from *T. annualata*-infected macrophages treated or not with 1μM or 5μM of the penetrating JNK-binding motif peptide and treated or not with MG132 (P), mutant (S>A) peptide (mP)or irrelevant peptide. (A) JNK2-IP shows western blot of the JNK2 precipitate probed with the anti-p104, anti-JNK2, and anti-GAPDH antibodies. Lower panel shows western blot analysis of immunoprecipitations probed with anti-JNK2 and anti-GAPDH antibodies. (1): V; (2): V+P 1μM; (3): V+P 5μM; (4): V+P 1μM + MG132; (5): V+P 5μM+ MG132; (6): V+ mP 1μM; (7): V+ mP 5μM; (8): IgG. (B) JNK1-IP: Immunoprecipitation analyses with anti-JNK1 using whole cell lysates derived from *T. annulata*-infected macrophages treated or not with 1 μM or 5μM of P or mP peptides. JNK1 protein expression was decreased following the treatment with JNK binding motif competitive peptide, while no effect was observed with the mP peptide. Lower panel shows western blot analysis of immunoprecipitations probed with anti-JNK1 and anti-GAPDH antibodies. (1): V; (2): V+P 1μM; (3): V+P 5μM; (4): V+mP 1μM; (5): V+mP 5μM; (6): IgG.

### Phosphorylation of the JNK-binding motif increases the affinity of p104 for JNK2

To understand the consequences of JNK2 association with p104, we specifically ablated their interaction. Located on the macroschizont surface GPI-anchored p104 has been described as being phosphorylated *in vivo* at several sites [30]. We noticed that two phosphorylated residues occurred in the conserved JNK-binding motif, specifically phospho-S806 and -S808 (TA08425 numbering). Consequently, penetrating peptides harbouring the conserved p104 decapeptide JNK-binding motif and a mutant peptide, where S806 and S808 had been replaced by alanine, together with an irrelevant peptide were synthesized (Table 1).

**Table 1.**
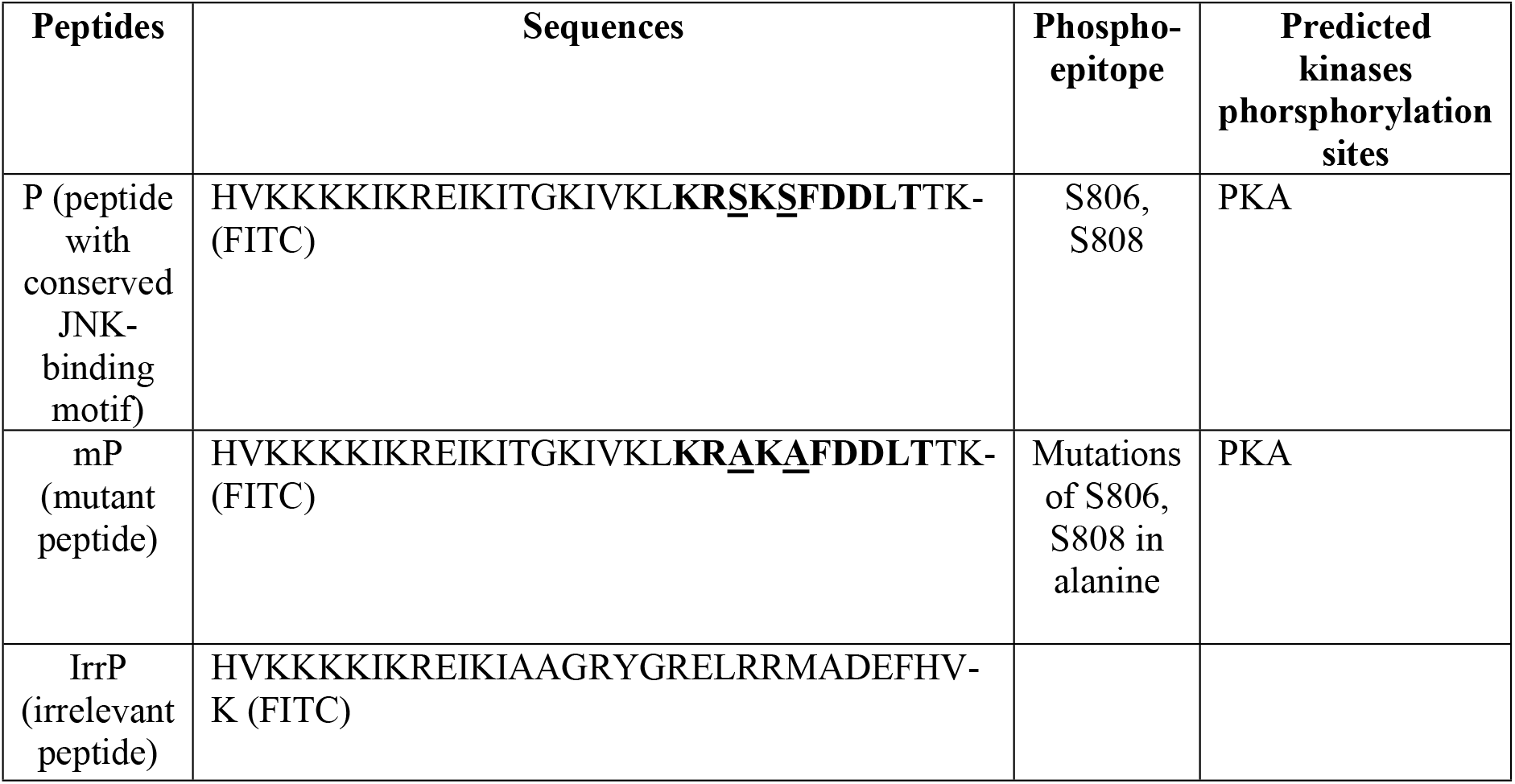
Synthesized peptides harbouring JNK binding motif. The different FITC-labelled penetrating peptides synthesized. (P) is the conserved wild type amino acid sequence with S806 and S808 shown underlined. The mutant peptide (mP) has S806 and S808 changed to A806 and A808 (underlined). An irrelevant peptide (IrrP) used as a negative control to compete for JNK-binding to p104. The JNK-binding motif is in bold and the sequence of the fused penetrating peptide indicated at the N-terminus.

All peptides were FITC conjugated and penetration into *T. annulata*-infected macrophages confirmed by immunofluorescence (FigS1). The peptide (P) corresponding to the wild type JNK-binding motif was able to ablate in dose-dependent manner JNK2 association with p104 (Figure 3). Importantly, the mutant (S>A) peptide (mP) did not abrogate JNK2 binding to p104 (Fig 3A, tracks 6 … 7), consistent with phosphorylation of S806 and/or S808 promoting p104 binding to JNK2. Following peptide-mediated abrogation of the JNK2/p104 complex the level of p54 JNK2 decreased, due to ubiquination of JNK2, but no drop in JNK2 levels was observed following proteasome blockade by MG132 (Fig 3A tracks 4 … 5 and Fig 4A). Although peptide-induced complex disruption reduced JNK2 levels no effect was observed on cytosolic JNK1 (Fig 4B), but loss of cytosolic JNK2 led to an increase in the amount of nuclear JNK1 (Fig 4B).

**Fig. 4.**
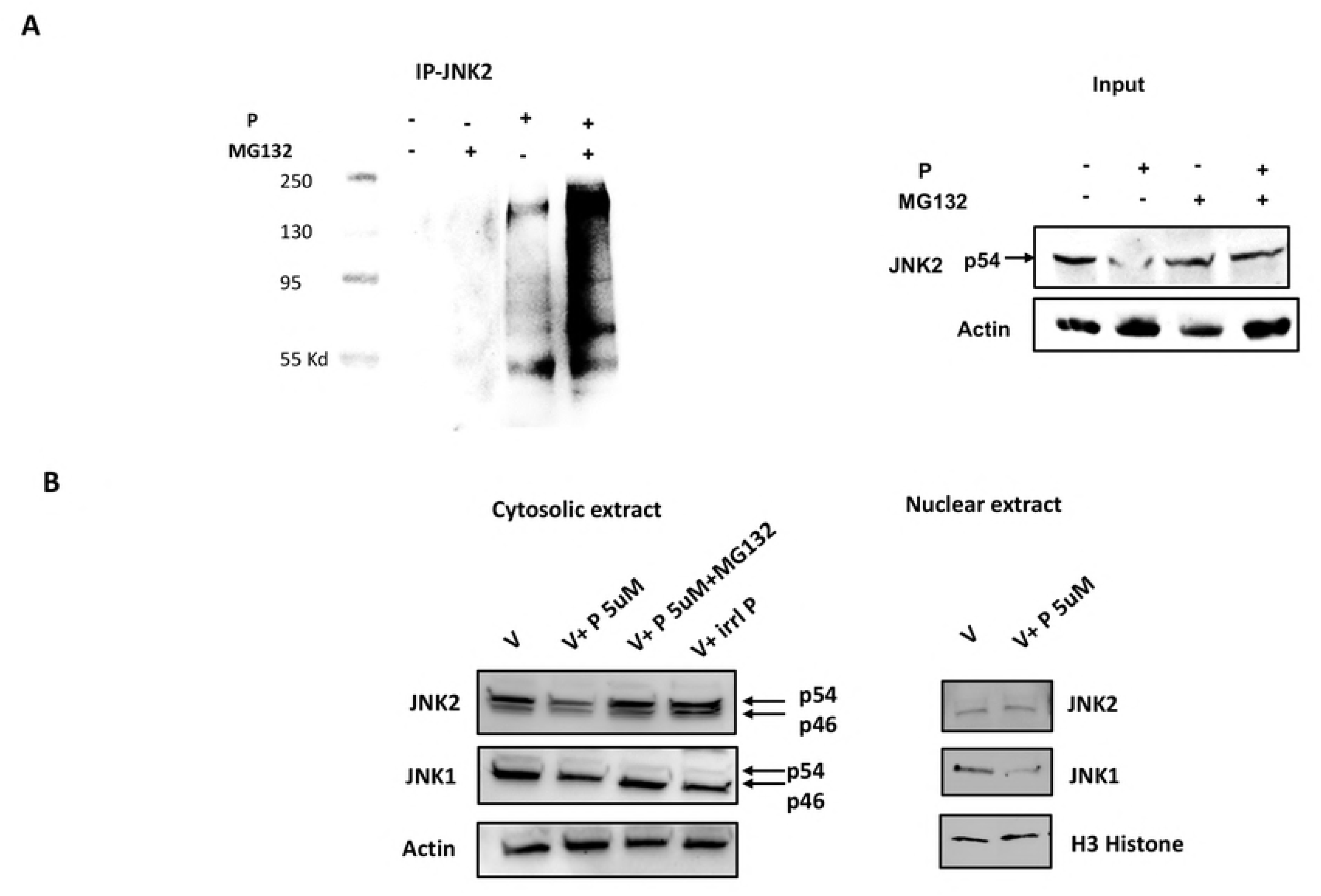
Association with p104 protects JNK2 from ubiquitination and proteosomal degradation. (A) Immunoprecipitation analyses with anti-JNK2 antibodies using whole cell lysates derived from virulent *T. annulata*-infected macrophages treated or not with 5μM of JNK-binding motif peptide P and treated or not with MG132. Western blot of the JNK2 precipitate probed with an anti-ubiquitin and JNK2 antibodies. (B). Western blot analysis of nuclear and cytosolic extracts probed with specific anti-JNK1 and anti-JNK2 antibodies using whole cell lysates derived from *T. annulata*-infected macrophages treated or not with 5μM of P and treated or not with MG132 and virulent treated with irrelevant peptide. Actin and H3 Histone antibodies were used as loading control. Input Panel A, JNK2 levels in the extracts were estimated compared to actin levels were used as loadingcontrol. Panel B, JNK1 and JNK2 levels in nuclear extracts were compared to histone H3 levels.

### PKA, but not JNK kinase activity contributes to p104 association with JNK2

Both S806 and S808 in p104 occur in a context (KRS*KS*FD) consistent with them being potentially phosphorylated by PKA [28]. Consequently, *T. annulata*-infected macrophages were treated for 2 h with myristoylated PKI or the ATP analogue H89 and both treatments dampened the association of p104 with JNK2 (Figure 5A). By contrast, treatment with a pan JNK inhibitor (pan-JNKi), or a JNK2-specific inhibitor (JNK2i) did not alter the ability of p104 to bind JNK2 (Fig 5B). We conclude that cAMP-dependent PKA likely phosphorylates S806 and/or S808 and their phosphorylation favours binding of p104 to bind JNK2. The kinase activity of JNK2 does not appear necessary for complex formation, leaving open that JNK2 could have a scaffold function (see below).

**Fig. 5.**
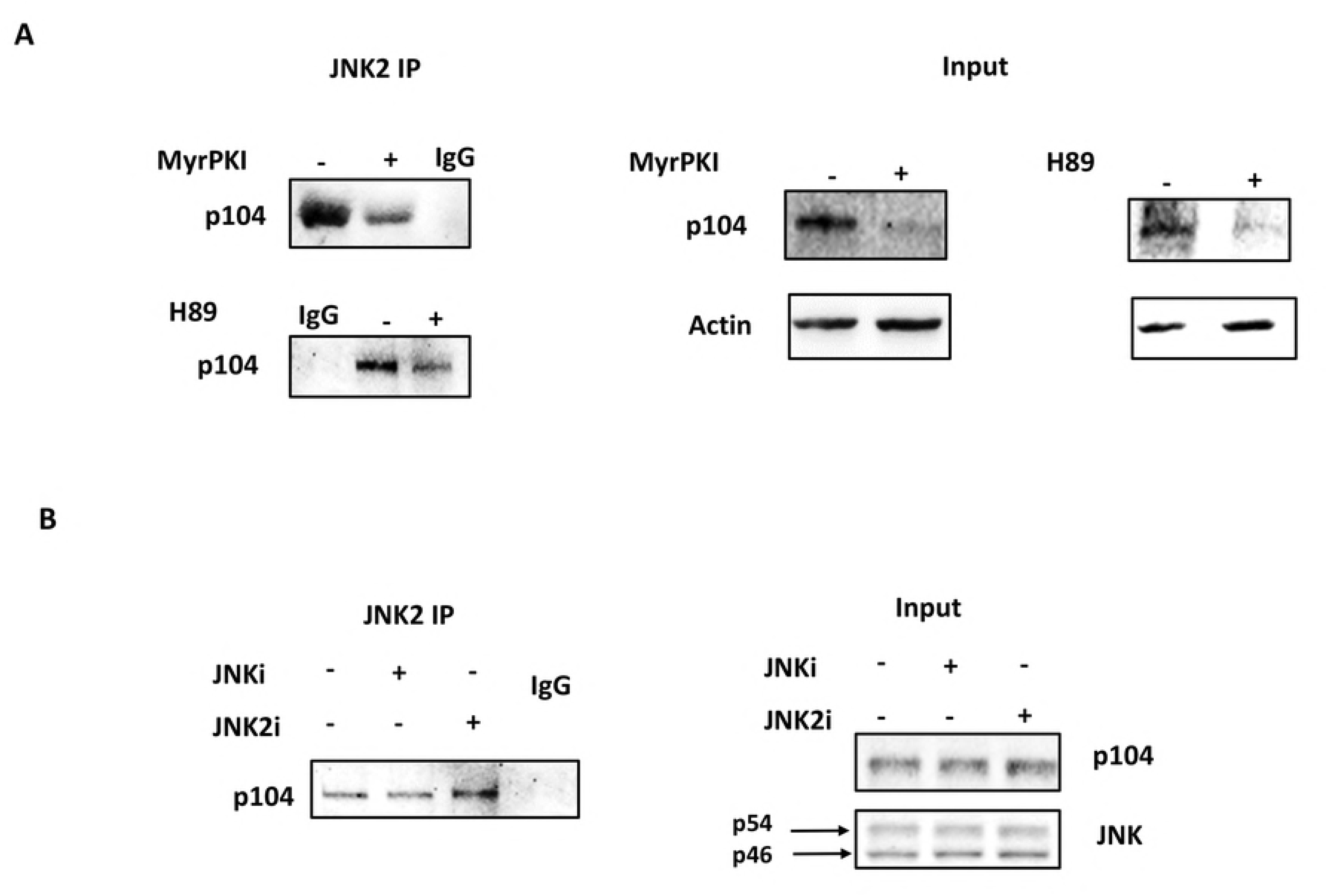
PKA phosphorylation increases association of p104 with JNK2. (A) Immunoprecipitation analyses with an anti-JNK2 antibody using whole cell lysates derived from virulent *T. annulata*-infected macrophages treated or not with the PKA specific inhibitor myristoylated PKI (MyrPKI) and H89. Left panel shows western blot of the JNK2 precipitate from non-treated, or MyrPKI/H89 treated (+) cells probed with the anti-p104 1C12 monoclonal antibody. Middle and right panels show the input levels of p104 and actin revealed by their respective antibodies. (B) Left panel: Western blot of JNK2 precipitates using extracts of cells treated with a pan-JNK inhibitor (pan-JNKi), or a JNK2-specific inhibitor (JNK2i) probed with the anti-p104 1C12 monoclonal antibody. Right panel: shows the input levels of p104 and the two JNK isoforms revealed by their respective antibodies.

### Abrogation of JNK2/p104 association reduces matrigel traversal of *Theileria-* transformed macrophages

Matrigel traversal is used as a measure of dissemination potential and virulent (V) *T. annulata*-transformed macrophages traverse matrigel better than attenuated (A) macrophages [20]. Matrigel transversal is significantly decreased when *T. annulata*-transformed macrophages are treated with the wild type JNK-binding motif peptide (P), whereas the (S>A) mutant peptide mP and the irrelevant peptide (irrP) had no significant effect (Figure 6A). It is well established that Theileria-infected macrophages are characterised by AP-1-driven transcription of *mmp9* and increased MMP9 activity promotes matrigel traversal and dissemination [21, 31, 32]. JNK2 association with p104 was therefore ablated and loss of MMP9 activity revealed by gelatin gel assay (Fig 6B, left). Peptide-induced disruption of JNK2/p104 binding slightly, but significantly, inhibited *mmp9* transcription as estimated by qRT-PCR (Fig 6B, right). Peptide-induced complex dissociation also specifically reduced nuclear c-Jun phosphorylation (Fig 6C) consistent with the drop in nuclear JNK1 levels (Fig 4B).

**Fig. 6.**
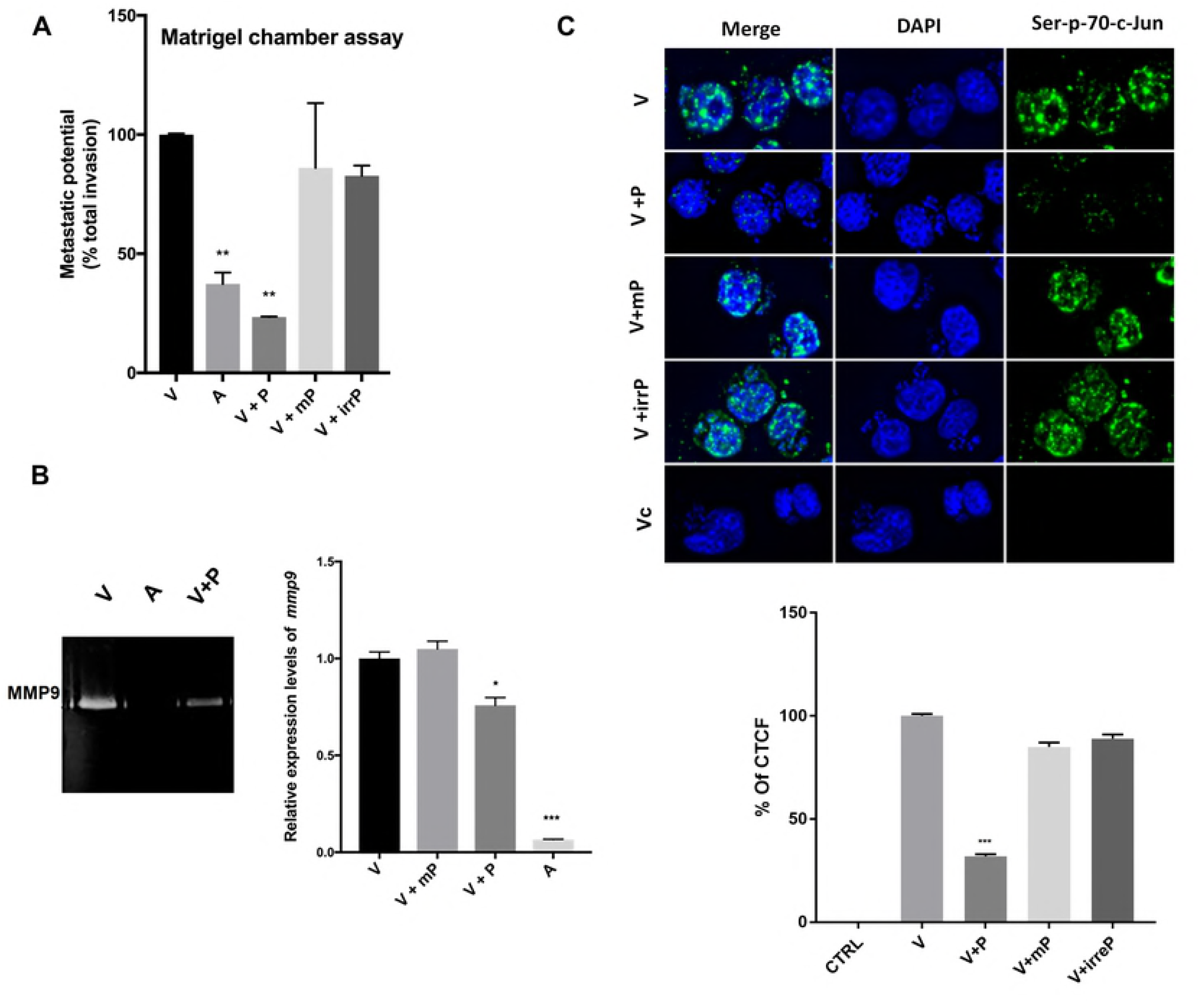
Peptide-provoked disruption of the JNK2/p104 complex diminishes matrigel traversal of *T. annulata*-transformed macrophages. (A) Upper panel: Matrigel traversal of virulent (V) compared to attenuated (A) macrophages and virulent macrophages treated with the JNK-motif penetrating peptide, the mutant (S>A) peptide (mP), or control irrelevant peptides (irrP). Peptide (P)-provoked disruption of the JNK2/p104 complex reduced matrigel traversal of virulent macrophages (V+P) to below attenuated levels, whereas treatment of virulent macrophages with the mutant (S>A) peptide (V+mP), or control peptide (irrP) had no effect. (B) Left panel. Zymogram showing MMP9 activity in the supernatants of virulent (V) compared to attenuated (A) macrophages and virulent macrophages treated with the competitive JNK2-binding peptide (P). Right panel. Relative expression of *mmp9* in virulent and attenuated *Theileria*-infected treated or not with the competitive JNK-binding peptide (P), or the mutant (S>A) peptide (mP). (C) Upper panel. Nuclear c-Jun phosphorylation displayed by virulent macrophages treated or not with the competitive JNK-binding peptide (P), the mutant (S>A) peptide (mP) or control peptides (irrP). Scale bar is equivalent to 10μ meters. Bottom panel. Percentage of corrected total cell fluorescence due to phospho-Ser73- c-Jun staining based on 30-independent cell images. All experiments were done independently (n = 3). The error bars show SEM values from 3 biological replicates, * p value <0.05, *** p value <0.001.

### Abrogation of the JNK2/p104 association leads to upregulation of ARF levels

Because a non-kinase, scaffold protein function for JNK2 has been described to regulate smARF levels and induce autophagy [33], we monitored ARF levels following disruption of the p104/JNK2 complex and subsequent JNK2 degradation (Figure 7). Loss of JNK2 provoked by peptide-treatment resulted in upregulation of p14ARF. The rise in ARF levels was accompanied an increase in amounts of processed autophagosome membrane protein LC3B-II and appearance of LC3B-II-positive foci (Figs 7A low panel, and 7B).

**Fig 7.**
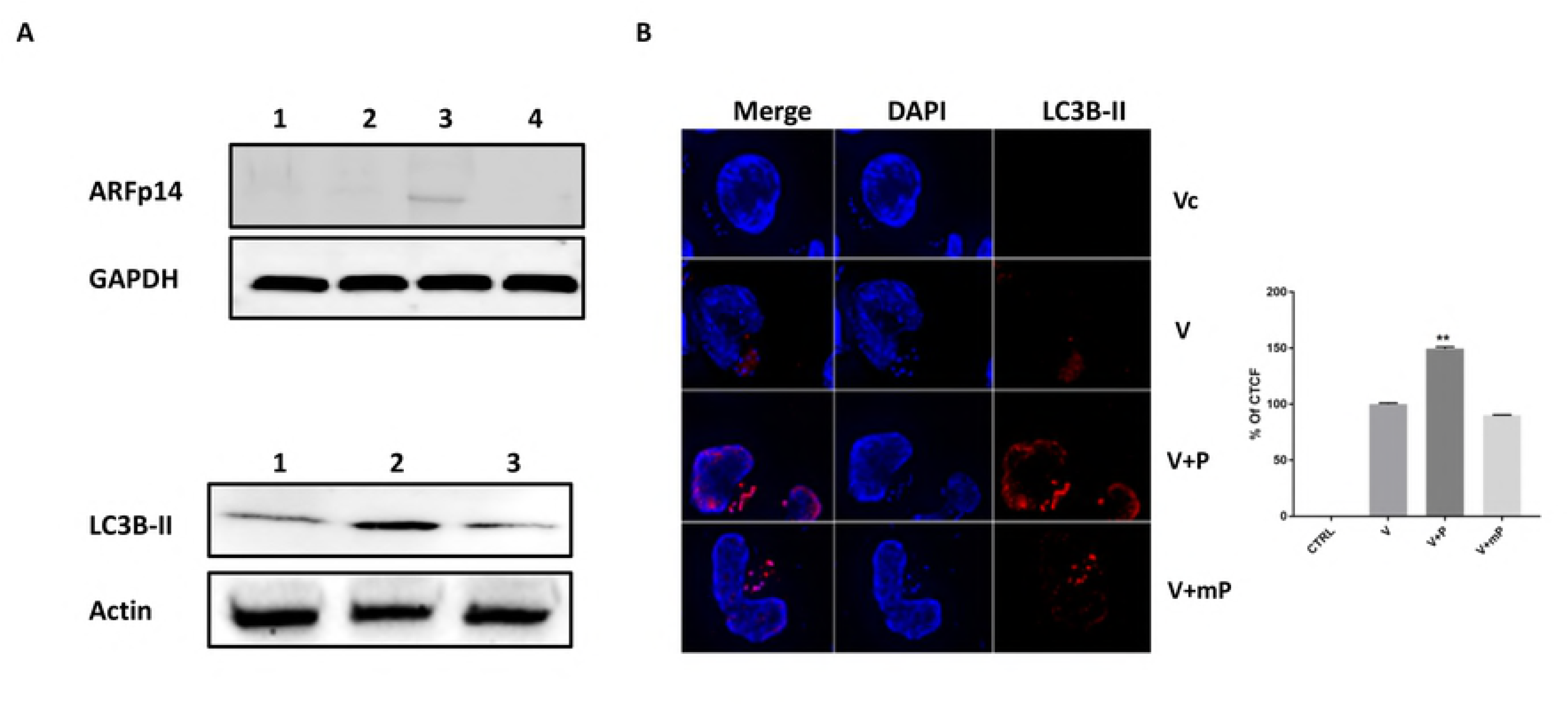
Loss of JNK2 provokes appearance of smARF and induction of autophagy. (A)Loss of JNK2 provoked by treating virulent macrophages (lane 1) with 1 μM (lane 2) and 5μM (lane 3) of the penetrating JNK-binding motif peptide causes a dose-dependent increase in the amounts of p14 ARF. No effect was observed with 5μM of mutant peptide (lane 4). Bottom, virulent macrophages (lane 1) were treated or not with 5μM of penetrating JNK-binding motif peptide (lane 2) and cell extracts probed with the specific LC3B-II antibody. 5μM peptide treatment resultes in augmented amounts of processed LC3B-II. No effect was observed with 5μM of mP (lane 3). (B). Immuofluorescence images obtained with anti-LC3B-II antibody using virulent (V) macrophages treated or not with JNK-binding motif peptide (P), or mutant peptide (mP). Only in peptide treated (V+P) macrophages is an augmentation in LC3B-II and clustering of LC3B-II-positive structures evident. No fluorescence was observed with Alexa-labelled secondary antibody (Vc). Scale bar is equivalent to 10μ meters. Bottom. Percentage of corrected total cell fluorescence due to LC3B-II staining based on 25- independent cell images. All experiments were done independently (n = 3). The error bars show SEM values from 3 biological replicates ** p value < 0.01.

## Discussion

Constitutively active JNK1 has a nuclear localisation that is dependent on live *T. annulata* macroschizonts, since nuclear JNK1 levels were ablated upon drug (Bw720c)-induced parasite death. By contrast, JNK2 is mainly in the infected macrophage cytosol associated with the macroschizont with only a minor fraction of JNK2 being nuclear and cytosolic JNK2 levels also depend on the live parasite being present (Figure 1). Simply by their different subcellular localisations one can surmise that JNK1 and JNK2 likely play different roles in *Theileria*-induced transformation of host leukocytes.

The close association between JNK2 and the parasite revealed by immunofluorescence led us to search for a parasite encoded JNK-binding protein located on the macroschizont surface. Interrogating the predicted *Theileria* proteomes with a consensus D-motif revealed the presence of 3 potential JNK-binding motifs in the GPI-anchored macroschizont *T. annulata* protein known as p104 [8]. We decided to characterise the putative JNK-binding site located between amino acids 804 to 814, as this is conserved in the *T. parva* p104 orthlogue (TP04_0437), and absent in the non-transforming *T. orientalis* (TOT_040000478) protein. Although not characterised here it remains possible that one or both of the other 2 *T. annulata* p104-specific D-motifs might also contribute to species-specific JNK2 binding by p104. *In vitro* recombinant *T. annulata* p104 clearly binds to recombinant JNK2 (FigS2A) making it possible to study the contribution of the 2 other D-motifs to species-specific JNK2 binding.

The conserved JNK-binding motif in p104 is distal to the previously described EB1-binding motif (SKIP) that’s located between amino acids 566 and 569 [28]. It is highly unlikely therefore that the JNK-binding motif penetrating peptide competes for EB1-binding. Furthermore, EB1 co-localizes with p104 on the macroschizont surface in a cell-cycle dependent manner, being more pronounced during cell division [28]. This contrasts with JNK2-binding to p104 that does not require division of infected macrophages. Although p104 acts as an EB1-binding protein attempts to interfere with EB1 binding to the macroschizont surface failed [28]. One explanation could be that JNK2 binding to p104 creates a platform favourable to EB1 association with the macroschizont during host cell mitosis. As we have shown that PKA can phosphorylate *in vitro* p104 (Fig S2B) this strongly argues that *in vivo* [0] PKA-mediated phosphorylation of p104 promotes association with JNK2 and indeed PKA inhibition reduced the amount of p104 detected in JNK2 immunoprecipitates (Fig 5). This implies that the complex at the surface of the macroschizont contains not only p104 and JNK2, but also PKA. By contrast, Cdk1-mediated phosphorylation of p104 seems to play a role in EB1 binding during mitosis, although the role of other kinases in regulating the interaction between p104 and EB1 has not been ruled out [28].

Importantly, both S806 and S808 have previously described as being phosphorylated *in vivo* [30], so the strategy we adopted to elucidate the role of the JNK2/p104 complex was to use a penetrating peptide harbouring the conserved JNK-binding motif, as a competitive p104 substrate for PKA. When S806 and S808 are changed to A806 and A808 the mutant penetrating peptide is no longer a competitive substrate for PKA and is incapable of disrupting JNK2 binding to p104. It’s noteworthy that binding of JNK2 to p104 *in vivo* was disrupted by inhibition of PKA activity by two independents PKA inhibitors, myristolated PKI and H89. Interestingly however, recruitment of JNK2 to p104 was insensitive to inhibition of JNK kinase activity suggesting JNK2 acts as a scaffold protein on which the complex is assembled at the macroschizont surface.

We focused on JNK2 to gain insights as to what might be the physiological advantage to *Theileria*-induced leukocyte transformation in retaining JNK2 in the infected host cell cytosol sequestered at the macroschizont surface. The ensembles of our results pinpoint at least 3 advantageous consequences of p104-mediated JNK2 sequestration: 1) While associated with the macroschizont surface JNK2 is protected from ubiquitination and proteasomal degradation and sustained JNK2 levels suppress smARF-mediated autophagy [11, 12]. As such JNK2 sequestration contributes to *Theileria*-infected leukocyte survival and indeed, 24 h following peptide-induced JNK2 degradation infected macrophages become Annexin V-positive (Fig S3). 2) Since nuclear JNK2 decreases c-Jun stability by promoting its ubiquitination [9, 10], cytosolic retention of JNK2 could contribute to sustained nuclear c-Jun levels perhaps by preventing DET1 mediated ubiquitination [34]. 3) Now, we show that peptide-induced complex dissociation led to JNK2 ubiquitination and degradation and a loss of nuclear c-Jun fluorescence (Fig 6C). Moreover, upon loss of cytosolic JNK2 the nuclear levels of JNK1 also decrease suggesting an alternative reason for loss of c-Jun phosphorylation (Fig 4B). Dampening of nuclear JNK1 levels also likely explains the drop in *mmp9* transcription and reduced matrigel traversal (Fig 6).

The macroschizont surface of *T. annulata*-infected macrophages is known to recruit another host cell tumour suppressor p53, preventing its translocation to the nucleus, inhibiting p53- mediated apoptosis, and thus contributing to host cell survival [35]. Moreover, ARF participates in the regulation of p53 interaction with MDM2 [36-38]. It’s interesting to note that antisense knockdown of *jnk2* has been described to dampen phosphorylation of p53 [39] making it possible that p53 is a substrate of JNK2 at the macroschizont surface in *Theileria-* infected leukocytes. It remains to be seen whether macroschizont recruitment of the IKK-signalosome [40] also involves binding to JNK2/p104, or since the number of parasite-associated IKK signalosomes fluctuates in the course of the host cell cycle, binding occurs indirectly perhaps via EB1, or other cytoskeleton-associated proteins.

Infection by another apicomplexan parasite *Toxoplasma gondii* leads to constitutive activation of a MAP/SAP kinase called p38 [41]. This contrasts with *Theileria,* where in different types of leukocytes transformed either by *T. annulata* or *T. parva* infection leads to constitutive activation of JNK rather than p38 [22, 42, 43]. Just why JNK2, and not JNK1 binds to p104 *in vivo* is not clear, since the JNK-binding motifs identified in p104 do not in principal discriminate between JNK isoforms. One possibility is that *in vivo* PKA-mediated phosphorylation of the conserved D-motif renders it more specific for JNK2 over JNK1. The *T. gondii* p38-binding protein harbours 2 MAP-kinase binding motifs, called KIM1 and KIM2 for Kinase-Interaction-Motifs (also known as D-motifs) that occur in a disordered C-terminal repetitive region of GR24 [44]. Although the MAP-kinase binding sites in p104 and GRA24 fit the same loose D-motif consensus their amino acid sequences are different. The 2 D-motifs present in GRA24 combine to provoke high affinity binding of p38, whereas we posit that PKA-mediated phosphorylation of S806 … S808 in p104 promotes binding to JNK2. It’s remarkable that these two pathogenic *Apicomplexa* both manipulate host cell MAP/SAP kinase signalling, but do so in different ways. Secreted GR24 goes into the host cell nucleus, binds and activates p38, whereas GPI-anchored p104 expressed on the macroschizont surface binds JNK2 preventing it from translocating to the nucleus, while activated JNK1 goes to the nucleus and phosphorylates c-Jun to drive *mmp9* transcription. Clearly, the need for a better understanding of both kinase and non-kinase, scaffold-like functions of JNK2 bound to p104 will animate future studies aimed at dissecting *Theileria*-induced leukocyte transformation.

## Materials and methods

### Chemicals and reagents

Pan-JNK inhibitor (JNK II #420128, Calbiochem, LA JOLLA) was added at 16 μM and JNK2 inhibitor (JNK IX: #420136, Calbiochem LA JOLLA) was added at 50 nM. Synthetized penetrating peptides harbouring JNK binding domain were produced by GL Biochem Ltd (Shanghai, China) and was added at 1 μM or 5μM for 2 h. MG132 (CAS 1211877-36-9) was added at 10 μM for 2 h. PKA inhibitor H89 (SIGMA-Aldrich) was added at 10 μM for 2 h and Myr PKI inhibitor (SIGMA-Aldrich) added at 50μM for 2 h.

### *Theileria annulata*-infected macrophage culture

*T. annulata*-infected monocytes/macrophages used in this study are the Ode virulent corresponding to passage 53 [45]. All cells were incubated at 37°C with 5% CO_2_ in Roswell Park Memorial Institute medium (RPMI) supplemented with 10% Fetal Bovine Serum (FBS), 2 mM L-Glutamine, 100 U penicillin, 0.1 mg/ml streptomycin, and 4-(2-hydroxyethyl)-1-piperazineethanesulfonic acid (HEPES).

### Analyses of JNK binding sites in *Theileria* proteomes

We obtained the protein sequences of *Theileria annulata, T. parva* and *T. orientalis* from PiroplasmDB. We used ‘dfinder’ programme, with default settings [46] to scan the complete predicted proteomes of the three species in search of D-sites (docking site for JNK). The search was filtered with cutoff threshold of 1e^-23^ as recommended [46]. Signal Peptide prediction was done by TOPCONS [47].

### Antibodies and western blot analyses

Cells were harvested and lysed by using lysis buffer (20 mM HEPES, 1% Nonidet P-40 [NP-40], 0.1% SDS, 150 mM NaCl, 2 mM EDTA, phosphatase inhibitor cocktail tablet (PhosSTOP; Roche), and protease inhibitor cocktail tablet (Complete Mini EDTA free; Roche). The protein concentration was determined by the Bradford protein assay. Cell lysates were subjected to Western blot analysis using conventional SDS-PAGE and protein transfer onto nitrocellulose filters (Protran and Whatman). Western blotting was performed as described previously [20]. The membrane was blocked with a solution containing 5% of BSA and Tris-buffered saline-Tween (TBST) for 1 h. The anti-*T. annulata* antibodies used and diluted in the blocking solution were the 1C12 monoclonal antibody against p104 [48] and an antibody to ribonucleotide reductase [49]. Polyclonal anti-JNK (sc-571), polyclonal anti-JNK2 (sc-46013), monoclonal anti-JNK2 (sc-271133), monoclonal anti-ubiquitine (sc-271289), polyclonal anti-phospho-c-Jun (sc-7981) and a polyclonal anti-p14^ARF^ (sc-8340) were purchased from Santa Cruz Biotechnologies (Santa Cruz, CA, USA), anti-LC3B-II (NB600-1384) from Novus Biologicals, the anti-PARP antibody (ab194586) from Abcam (Abcam PLC, Cambridge) and anti-MMP9 (AV33090) from Sigma. After washing, proteins were visualized with ECL western blotting detection reagents (Thermo Scientific) on fusion FX (Vilber Lourmat). The β-actin level was used as a loading control throughout.

### Immunofluorescence analysis

Ode macrophages were treated or not with peptide and fixed in buffer containing 4% paraformaldehyde and 3% of sucrose for 15 min. Permeabilization was performed using 0.01% Triton in phosphate-buffered saline medium for 5 min followed by two washes with 1X phosphate-buffered saline. Cells were then blocked with 3% bovine serum albumin for 1 h, stained with the anti-LC3B-II antibody, or the anti-phospho-c-Jun for 120 min at room temperature, and washed 3 times with buffer before incubation with a secondary anti-rabbit IgG antibody conjugated with respectively Alexa 546 or Alexa 647 (Molecular Probes) in darkness for 60 min at room temperature. Cells were stained with 4’,6- diamidino-2-phenylindole (Bibenzimide H 33258, Sigma) for nucleus labeling. Dako mounting medium was used (Glostrup, Denmark). The immunolabeled cells were examined with a Zeiss observer Z1, camera QICAM.

### Co-immunoprecipitation

*T. annulata*-infected macrophages were harvested and lysed in the lysis buffer containing 20 mM HEPES, 1% Nonidet P-40 [NP-40], 0.1% SDS, 150 mM NaCl, 2 mM EDTA, phosphatase inhibitor cocktail tablet (PhosSTOP; Roche), and protease inhibitor cocktail tablet (Complete Mini EDTA free; Roche). The protein concentration was determined by the Bradford protein assay. Protein-G Dynabeads (Invitrogen) were washed twice with PBS1X solution. After incubation with the antibody of interest for 2.5 h, 500 μg of protein extracts were added overnight. The beads were washed 5-times with lysis buffer supplemented with proteases and phosphatases inhibitors and boiled in Laemmli buffer before performing western blotting.

### Matrigel chambers assay

The invasive capacity of *Theileria*-infected macrophages was assessed *in vitro* using matrigel migration chambers [16]. Culture coat 96-well medium BME cell invasion assay was obtained from Culturex instructions (3482-096-K). After 24 h of incubation at 37°C, each well of the top chamber was washed once in buffer. The top chamber was placed back on the receiver plate. 100μl of cell dissociation solution/Calcein AM were added to the bottom chamber of each well, incubated at 37°C for 1 h to fluorescently label cells and dissociate them from the membrane before reading at 485 nm excitation, 520 nm emission using the same parameters as the standard curve.

### Zymography (gelatin gel assay)

We used 10% SDS-Polyacrylamide gel electrophoresis containing 1% co-polymerized gelatin to detect secreted gelatinases such as MMP2 and MMP9. 5×106 cells were washed three times with cold PBS to remove all the serum and were plated in 6-well plates in 5ml serum free culture medium. Supernatants from these cultures were collected after 24 h. Supernatant samples were mixed with 2X sample buffer containing 0.5M Tris-HCl pH 6.8, 20% Glycerol, 10% SDS and 0.005% Bromophenol Blue. They were left at room temperature for 10 min and then loaded onto the gel. Migration was performed in 1X Tris-Glycine SDS Running buffer at 125V. The gels were washed twice for 30 min in renaturing buffer containing 2.5% Triton X-100, which removed the SDS. To activate the proteases, gels were incubated at 37°C for 18 h in 30ml of a solution containing 50mM Tris-HCl pH 7.6, 5mM CaCl2 and 0.02% Triton X-100. Gels were subsequently stained for 2 h with a solution containing 0.5% Coomasie Blue R-250, 40% Methanol, 10% Acetic acid and de-stained with 50% Methanol, 10% Acetic Acid. Areas of digestion appeared as clear bands, against a darkly stained background due to the substrate being degraded by the enzyme.

### RNA extraction, reverse transcription and qRT-PCR

Total RNA was extracted from cells with the RNeasy^®^ Plus mini kit (QIAGEN) and quantified by the NanoDrop ND1000 Spectrophotometer. cDNA was synthesized from 1000 ng of RNA by using M-MLV reverse transcriptase enzyme (Promega). The qRT-PCR reaction mixture included 2.5 μl Absolute blue qPCR SYBR green (Thermo Scientific), 0.5 μl of each forward and reverse primers, 4 μl molecular grade water, and 2.5 μl of 1:20 diluted cDNA.

Actin left primer: AGAGGCATCCTGACCCTCAA
Actin right primer: TCTCCATGTCGTCCCAGTTG
MMP9 left: TGGCACGGAGGTGTGATCTA
MMP9 right: GACAAGAAGTGGGGCTTCTG

### GST-pull downs

<u>The C-terminal disordered region (504-839) of</u> *T. annulata* p104 was subcloned into a modified pET vector enabling expression of GST fusion proteins with a C-terminal hexa-histidine tag by PCR. The cDNA of full-length human JNK2 was subcloned into another modified pET plasmid allowing the expression of proteins with an N-terminal hexahistidine tag. All protein constructs were expressed in *Escherichia coli* Rosetta (DE) pLysS cells with standard techniques. Protein expression was induced at 25 °C for 3 h by adding 0.2 mM IPTG, cells were lysed, and the lysate was loaded onto Ni-NTA resin and eluted by imidazol. GST-p104 samples were then loaded to glutathione resin and washed with GST wash buffer (20 mM Tris pH 8.0, 150 mM NaCl, 0.05 % IGEPAL, 1mM EDTA and 5 mM beta-mercaptoethanol). Ni-NTA eluted JNK2 was further purified by using an ionexchange column (resourceQ) and was eluted with a salt gradient (0.1M-1M NaCl). In a typical GST-pull down experiment 50 μl of glutathion resin loaded with the bait was incubated in 100 μM of JNK2 solution in GST wash buffer and washed three times. After addition of SDS loading buffer the resin was subjected to SDS-PAGE and gels were stained by Coomassie Brilliant Blue protein dye, or the gels were subjected to Western-blots using anti-His, GE Healthcare (27-4710-01), or anti-JNK, Cell Signaling (3708S), antibodies according to the supplier’s recommendations. All plasmid DNA sequences were confirmed by sequencing. GST protein with a C-terminal hexa-histidine tag was used as the negative control for the GST-pull down experiments.

### *In vitro* kinase assays

The catalytic domain of PKA with an N-terminal hexa-histidine tag was expressed in *E. coli* using the pET15b PKA Cat vector [0] and purified with Ni-NTA resin similarly as described above. 0.5 μM PKA catalytic subunit was incubated with 5μM GST or GST-p104 C-terminal disordered region (504-83 9) fusion protein in the presence of radioactively labeled ATP(γ)P^32^ (~5 μCi). Aliquots of the kinase reactions were taken at different time points and run on SDS-PAGE. Gels were mounted onto filter paper, dried and subjected to phosphoimaging using a Typhoon Trio+ scanner (GE Healthcare). The kinase buffer contained 20 mM Tris pH, 100 mM NaCl, 0.05 % IGEPAL, 5 % glycerol, 2 mM TCEP, 5 mM MgCl_2_ and 0.25 mM ATP.

### Flow cytometry

Infected macrophages were treated with 5 μM of non-constrained CCP. 10^6^ of Ode cells are prepared in 1 ml of PBS with 10% FBS in each test tube. After a centrifugation for 5 min at 200×g and 4 °C, cells are resuspended in 100 μl annexin V Binding buffer. 5 μl of annexin V and 5 μl of 7AAD (7-aminoactinomycin D) are added to each tube except single stained control. Ode are incubated 15 min in the dark at room temperature with 400 μl ice cold annexin V binding buffer and then analyzed on the flow cytometry (Accuri C6 – C flow Plus software).

### Statistical analysis

Experiments were performed at least three times and results presented as mean values +/- SEM. p values were determined using the Student’s t-test. Results were considered significant for p <0.05.

## Acknowledgments

We would like to thank Professor Brian Shiels for gift of antibodies to *Theileria* p104 (mAb 1C12).

## Supporting information

**S1 file:** Screening for potential JNK binding sites (D-site) using dfinder with a 1e-23 cutoff and the predicted proteomes of *Theileria annulata*, *T. parva* and *T. orientalis*.

**S1 Fig:** Penetrating peptides enter *T. annulata-infected* macrophages. Virulent *T*. annulata-infected macrophages are treated with FITC-conjugated peptide for 2h. Green: peptide; Dapi: nucleus.

**S2 Fig:** (A) GST pulldown with JNK2 and p104 recombinant proteins. Recombinant proteins were expressed in *E. coli* and purified. Baits (lane 1, GST+ JNK2, Lane 2, p104, Lane 3, JNK2) were loaded to glutathion resin and were incubated with JNK2 (Load, Lane 4). Samples were run on SDS-PAGE and the gel was stained with Coomassiee dye, or was subjected to Western blot and protein bands visualized by an anti-His antibody (as all purified proteins had a 6xHis tag). (M - Marker, molecular weights are shown in kDa at the left). **(B) *In vitro* PKA-mediated phosphorylation of recombinant p104.** 0.5 μM of the PKA catalytic subunit was incubated with 5μM GST or GST-p104 fusion proteins in the presence of radioactively labelled ATP(γ)_P_32. Aliquots of the kinase reactions were taken at the indicated time points and run on SDS-PAGE. The upper panel shows the gel stained with Coomassie dye and the lower panel shows the phosphor imaging results of the same gel. (M - Marker, molecular weights are shown in kDa at the left).

**S3 Figure :** 24 h after peptide-induced disruption of JNK2/p104 complex *Theileria-* infected macrophages become Annexin V-positive. Infected macrophages were treated for 24 h with 1 μM or 5 μM of JNK-motif penetrating peptide or the mutant (S>A) peptide (mP) in presence or absence of MG132 and the percentage of cells undergoing apoptosis was estimated by FACS using annexin V/7AAD staining.

## References

1. Davis RJ. Signal transduction by the JNK group of MAP kinases. Cell. 2000;103(2):239–52. PubMed PMID: 11057897.

2. Bogoyevitch MA, Kobe B. Uses for JNK: the many and varied substrates of the c-Jun N-terminal kinases. Microbiol Mol Biol Rev. 2006;70(4):1061–95. doi: 10.1128/MMBR.00025-06. PubMed PMID: 17158707; PubMed Central PMCID: PMCPMC1698509.

3. Hibi M, Lin A, Smeal T, Minden A, Karin M. Identification of an oncoprotein-and UV-responsive protein kinase that binds and potentiates the c-Jun activation domain. Genes Dev. 1993;7(11):2135–48. PubMed PMID: 8224842.

4. Nateri AS, Riera-Sans L, Da Costa C, Behrens A. The ubiquitin ligase SCFFbw7 antagonizes apoptotic JNK signaling. Science. 2004;303(5662):1374–8. doi: 10.1126/science.1092880. PubMed PMID: 14739463.

5. Gao M, Labuda T, Xia Y, Gallagher E, Fang D, Liu YC, et al. Jun turnover is controlled through JNK-dependent phosphorylation of the E3 ligase Itch. Science. 2004;306(5694):271–5. doi: 10.1126/science.1099414. PubMed PMID: 15358865.

6. Huang C, Rajfur Z, Borchers C, Schaller MD, Jacobson K. JNK phosphorylates paxillin and regulates cell migration. Nature. 2003;424(6945):219–23. doi: 10.1038/nature01745. PubMed PMID: 12853963.

7. Ueno H, Tomiyama A, Yamaguchi H, Uekita T, Shirakihara T, Nakashima K, et al. Augmentation of invadopodia formation in temozolomide-resistant or adopted glioma is regulated by c-Jun terminal kinase-paxillin axis. Biochem Biophys Res Commun. 2015;468(1-2):240–7. doi: 10.1016/j.bbrc.2015.10.122. PubMed PMID: 26518652.

8. Wang C, Zhao Y, Su Y, Li R, Lin Y, Zhou X, et al. C-Jun N-terminal kinase (JNK) mediates Wnt5a-induced cell motility dependent or independent of RhoA pathway in human dental papilla cells. PLoS One. 2013;8(7):e69440. doi: 10.1371/journal.pone.0069440. PubMed PMID: 23844260; PubMed Central PMCID: PMCPMC3700942.

9. Bode AM, Dong Z. The functional contrariety of JNK. Mol Carcinog. 2007;46(8):591–8. doi: 10.1002/mc.20348. PubMed PMID: 17538955; PubMed Central PMCID: PMCPMC2832829.

10. Fuchs SY, Dolan L, Davis RJ, Ronai Z. Phosphorylation-dependent targeting of c-Jun ubiquitination by Jun N-kinase. Oncogene. 1996;13(7):1531–5. PubMed PMID: 8875991.

11. Budina-Kolomets A, Hontz RD, Pimkina J, Murphy ME. A conserved domain in exon 2 coding for the human and murine ARF tumor suppressor protein is required for autophagy induction. Autophagy. 2013;9(10):1553–65. doi: 10.4161/auto.25831. PubMed PMID: 23939042; PubMed Central PMCID: PMCPMC4623555.

12. Zhang Q, Kuang H, Chen C, Yan J, Do-Umehara HC, Liu XY, et al. The kinase Jnk2 promotes stress-induced mitophagy by targeting the small mitochondrial form of the tumor suppressor ARF for degradation. Nat Immunol. 2015;16(5):458–66. doi: 10.1038/ni.3130. PubMed PMID: 25799126; PubMed Central PMCID: PMCPMC4451949.

13. Darghouth MA. Review on the experience with live attenuated vaccines against tropical theileriosis in Tunisia: considerations for the present and implications for the future. Vaccine. 2008;26 Suppl 6:G4–G10. doi: 10.1016/j.vaccine.2008.09.065. PubMed PMID: 19178892.

14. Baylis HA, Megson A, Hall R. Infection with Theileria annulata induces expression of matrix metalloproteinase 9 and transcription factor AP-1 in bovine leucocytes. Mol Biochem Parasitol. 1995;69(2):211–22. PubMed PMID: 7770085.

15. Fell AH, Preston PM, Ansell JD. Establishment of Theileria-infected bovine cell lines in scid mice. Parasite Immunol. 1990;12(3):335–9. PubMed PMID: 2117267.

16. Lizundia R, Chaussepied M, Huerre M, Werling D, Di Santo JP, Langsley G. c-Jun NH2-terminal kinase/c-Jun signaling promotes survival and metastasis of B lymphocytes transformed by Theileria. Cancer Res. 2006;66(12):6105–10. doi: 10.1158/0008-5472.CAN-05-3861. PubMed PMID: 16778183.

17. Shiels B, Langsley G, Weir W, Pain A, McKellar S, Dobbelaere D. Alteration of host cell phenotype by Theileria annulata and Theileria parva: mining for manipulators in the parasite genomes. Int J Parasitol. 2006;36(1):9–21. doi: 10.1016/j.ijpara.2005.09.002. PubMed PMID: 16221473.

18. Chaussepied M, Janski N, Baumgartner M, Lizundia R, Jensen K, Weir W, et al. TGF-b2 induction regulates invasiveness of Theileria-transformed leukocytes and disease susceptibility. PLoS Pathog. 2010;6(11):e1001197. doi: 10.1371/journal.ppat.1001197. PubMed PMID: 21124992; PubMed Central PMCID: PMCPMC2987823.

19. Haidar M, Echebli N, Ding Y, Kamau E, Langsley G. Transforming growth factor beta2 promotes transcription of COX2 and EP4, leading to a prostaglandin E2-driven autostimulatory loop that enhances virulence of Theileria annulata-transformed macrophages. Infect Immun. 2015;83(5):1869–80. doi: 10.1128/IAI.02975-14. PubMed PMID: 25690101; PubMed Central PMCID: PMCPMC4399038.

20. Haidar M, Whitworth J, Noe G, Liu WQ, Vidal M, Langsley G. TGF-beta2 induces Grb2 to recruit PI3-K to TGF-RII that activates JNK/AP-1-signaling and augments invasiveness of Theileria-transformed macrophages. Sci Rep. 2015;5:15688. doi: 10.1038/srep15688. PubMed PMID: 26511382; PubMed Central PMCID: PMCPMC4625156.

21. Adamson R, Logan M, Kinnaird J, Langsley G, Hall R. Loss of matrix metalloproteinase 9 activity in Theileria annulata-attenuated cells is at the transcriptional level and is associated with differentially expressed AP-1 species. Mol Biochem Parasitol. 2000;106(1):51–61. PubMed PMID: 10743610.

22. Chaussepied M, Lallemand D, Moreau MF, Adamson R, Hall R, Langsley G. Upregulation of Jun and Fos family members and permanent JNK activity lead to constitutive AP-1 activation in Theileria-transformed leukocytes. Mol Biochem Parasitol. 1998;94(2):215–26. PubMed PMID: 9747972.

23. Lizundia R, Chaussepied M, Naissant B, Masse GX, Quevillon E, Michel F, et al. The JNK/AP-1 pathway upregulates expression of the recycling endosome rab11a gene in B cells transformed by Theileria. Cell Microbiol. 2007;9(8):1936–45. doi: 10.1111/j.1462-5822.2007.00925.x. PubMed PMID: 17388783.

24. Dobbelaere D, Heussler V. Transformation of leukocytes by Theileria parva and T. annulata. Annu Rev Microbiol. 1999;53:1–42. doi: 10.1146/annurev.micro.53.1.1. PubMed PMID: 10547684.

25. Lizundia R, Sengmanivong L, Guergnon J, Muller T, Schnelle T, Langsley G, et al. Use of micro-rotation imaging to study JNK-mediated cell survival in Theileria parva-infected B-lymphocytes. Parasitology. 2005;130(Pt 6):629–35. PubMed PMID: 15977899.

26. Metheni M, Echebli N, Chaussepied M, Ransy C, Chereau C, Jensen K, et al. The level of H(2)O(2) type oxidative stress regulates virulence of Theileria-transformed leukocytes. Cell Microbiol. 2014;16(2):269–79. doi: 10.1111/cmi.12218. PubMed PMID: 24112286; PubMed Central PMCID: PMCPMC3906831.

27. Metheni M, Lombes A, Bouillaud F, Batteux F, Langsley G. HIF-1alpha induction, proliferation and glycolysis of Theileria-infected leukocytes. Cell Microbiol. 2015;17(4):467–72. doi: 10.1111/cmi.12421. PubMed PMID: 25620534.

28. Woods KL, Theiler R, Muhlemann M, Segiser A, Huber S, Ansari HR, et al. Recruitment of EB1, a master regulator of microtubule dynamics, to the surface of the Theileria annulata schizont. PLoS Pathog. 2013;9(5):e1003346. doi: 10.1371/journal.ppat.1003346. PubMed PMID: 23675298; PubMed Central PMCID: PMCPMC3649978.

29. Zeke A, Bastys T, Alexa A, Garai A, Meszaros B, Kirsch K, et al. Systematic discovery of linear binding motifs targeting an ancient protein interaction surface on MAP kinases. Mol Syst Biol. 2015;11(11):837. doi: 10.15252/msb.20156269. PubMed PMID: 26538579; PubMed Central PMCID: PMCPMC4670726.

30. Wiens O, Xia D, von Schubert C, Wastling JM, Dobbelaere DA, Heussler VT, et al. Cell cycle-dependent phosphorylation of Theileria annulata schizont surface proteins. PLoS One. 2014;9(7):e103821. doi: 10.1371/journal.pone.0103821. PubMed PMID: 25077614; PubMed Central PMCID: PMCPMC4117643.

31. Cock-Rada AM, Medjkane S, Janski N, Yousfi N, Perichon M, Chaussepied M, et al. SMYD3 promotes cancer invasion by epigenetic upregulation of the metalloproteinase MMP-9.Cancer Res. 2012;72(3):810–20. doi: 10.1158/0008-5472.CAN-11-1052. PubMed PMID: 22194464; PubMed Central PMCID: PMCPMC3299564.

32. Echebli N, Mhadhbi M, Chaussepied M, Vayssettes C, Di Santo JP, Darghouth MA, et al. Engineering attenuated virulence of a Theileria annulata-infected macrophage. PLoS Negl Trop Dis. 2014;8(11):e3183. doi: 10.1371/journal.pntd.0003183. PubMed PMID: 25375322; PubMed Central PMCID: PMCPMC4222746.

33. Zhang Q, Kuang H, Chen C, Yan J, Do-Umehara HC, Liu XY, et al. Corrigendum: The kinase Jnk2 promotes stress-induced mitophagy by targeting the small mitochondrial form of the tumor suppressor ARF for degradation. Nat Immunol. 2015;16(7):785. doi: 10.1038/ni0715-785b. PubMed PMID: 26086147.

34. Marsolier J, Pineau S, Medjkane S, Perichon M, Yin Q, Flemington E, et al. OncomiR addiction is generated by a miR-155 feedback loop in Theileria-transformed leukocytes. PLoS Pathog. 2013;9(4):e1003222. doi: 10.1371/journal.ppat.1003222. PubMed PMID: 23637592; PubMed Central PMCID: PMCPMC3630095.

35. Haller D, Mackiewicz M, Gerber S, Beyer D, Kullmann B, Schneider I, et al. Cytoplasmic sequestration of p53 promotes survival in leukocytes transformed by Theileria. Oncogene. 2010;29(21):3079–86. doi: 10.1038/onc.2010.61. PubMed PMID: 20208567.

36. Maggi LB, Jr., Winkeler CL, Miceli AP, Apicelli AJ, Brady SN, Kuchenreuther MJ, et al. ARF tumor suppression in the nucleolus. Biochim Biophys Acta. 2014;1842(6):831–9. doi: 10.1016/j.bbadis.2014.01.016. PubMed PMID: 24525025.

37. Trino S, De Luca L, Laurenzana I, Caivano A, Del Vecchio L, Martinelli G, et al. P53-MDM2 Pathway: Evidences for A New Targeted Therapeutic Approach in B-Acute Lymphoblastic Leukemia. Front Pharmacol. 2016;7:491. doi: 10.3389/fphar.2016.00491. PubMed PMID: 28018226; PubMed Central PMCID: PMCPMC5159974.

38. Vivo M, Matarese M, Sepe M, Di Martino R, Festa L, Calabro V, et al. MDM2-mediated degradation of p14ARF: a novel mechanism to control ARF levels in cancer cells. PLoS One. 2015;10(2):e0117252. doi: 10.1371/journal.pone.0117252. PubMed PMID: 25723571; PubMed Central PMCID: PMCPMC4344200.

39. Buschmann T, Potapova O, Bar-Shira A, Ivanov VN, Fuchs SY, Henderson S, et al. Jun NH2-terminal kinase phosphorylation of p53 on Thr-81 is important for p53 stabilization and transcriptional activities in response to stress. Mol Cell Biol. 2001;21(8):2743–54. doi:10.1128/MCB.21.8.2743-2754.2001. PubMed PMID: 11283254; PubMed Central PMCID: PMCPMC86905.

40. Heussler VT, Rottenberg S, Schwab R, Kuenzi P, Fernandez PC, McKellar S, et al. Hijacking of host cell IKK signalosomes by the transforming parasite Theileria. Science. 2002;298(5595):1033–6. doi: 10.1126/science.1075462. PubMed PMID: 12411708.

41. Braun L, Brenier-Pinchart MP, Yogavel M, Curt-Varesano A, Curt-Bertini RL, Hussain T, et al. A Toxoplasma dense granule protein, GRA24, modulates the early immune response to infection by promoting a direct and sustained host p38 MAPK activation. J Exp Med. 2013;210(10):2071–86. doi: 10.1084/jem.20130103. PubMed PMID: 24043761; PubMed Central PMCID: PMCPMC3782045.

42. Galley Y, Hagens G, Glaser I, Davis W, Eichhorn M, Dobbelaere D. Jun NH2-terminal kinase is constitutively activated in T cells transformed by the intracellular parasite Theileria parva. Proc Natl Acad Sci U S A. 1997;94(10):5119–24. PubMed PMID: 9144200; PubMed Central PMCID: PMCPMC24641.

43. Botteron C, Dobbelaere D. AP-1 and ATF-2 are constitutively activated via the JNK pathway in Theileria parva-transformed T-cells. Biochem Biophys Res Commun. 1998;246(2):418–21. doi: 10.1006/bbrc.1998.8635. PubMed PMID: 9610375.

44. Pellegrini E, Palencia A, Braun L, Kapp U, Bougdour A, Belrhali H, et al. Structural Basis for the Subversion of MAP Kinase Signaling by an Intrinsically Disordered Parasite Secreted Agonist. Structure. 2017;25(1):16–26. doi: 10.1016/j.str.2016.10.011. PubMed PMID: 27889209; PubMed Central PMCID: PMCPMC5222587.

45. Singh S, Khatri N, Manuja A, Sharma RD, Malhotra DV, Nichani AK. Impact of field vaccination with a Theileria annulata schizont cell culture vaccine on the epidemiology of tropical theileriosis. Vet Parasitol. 2001;101(2):91–100. PubMed PMID: 11587838.

46. Whisenant TC, Ho DT, Benz RW, Rogers JS, Kaake RM, Gordon EA, et al. Computational prediction and experimental verification of new MAP kinase docking sites and substrates including Gli transcription factors. PLoS Comput Biol. 2010;6(8). doi: 10.1371/journal.pcbi.1000908. PubMed PMID: 20865152; PubMed Central PMCID: PMCPMC2928751.

47. Tsirigos KD, Peters C, Shu N, Kall L, Elofsson A. The TOPCONS web server for consensus prediction of membrane protein topology and signal peptides. Nucleic Acids Res. 2015;43(W1):W401–7. doi: 10.1093/nar/gkv485. PubMed PMID: 25969446; PubMed Central PMCID: PMCPMC4489233.

48. Shiels BR, McDougall C, Tait A, Brown CG. Identification of infection-associated antigens in Theileria annulata transformed cells. Parasite Immunol. 1986;8(1):69–77. PubMed PMID: 2421227.

49. Swan DG, Stadler L, Okan E, Hoffs M, Katzer F, Kinnaird J, et al. TashHN, a Theileria annulata encoded protein transported to the host nucleus displays an association with attenuation of parasite differentiation. Cell Microbiol. 2003;5(12):947–56. PubMed PMID: 14641179.

50. Narayana N, Cox S, Shaltiel S, Taylor SS, Xuong N. Crystal structure of a polyhistidine-tagged recombinant catalytic subunit of cAMP-dependent protein kinase complexed with the peptide inhibitor PKI(5-24) and adenosine. Biochemistry. 1997;36(15):4438–48. doi: 10.1021/bi961947+. PubMed PMID: 9109651.

